# The (*α, β*)-*k* Boolean Signatures of Molecular Toxicity: Microcystin as a Case Study

**DOI:** 10.1101/2024.12.29.630644

**Authors:** Pablo Moscato, Sabrina Jaeger-Honz, Mohammad Nazmul Haque, Falk Schreiber

## Abstract

**Background:** The (*α, β*)-*k*-Feature Set Problem is a combinatorial problem, that has been proven as alternative to typical methods for reducing the dimensionality of large datasets without compromising the performance of machine learning classifiers.

**Result:** We present a case study that shows that solutions of the (*α, β*)-*k*-Feature Set Problem help to identify molecular substructures related to toxicity. The dataset investigated in this study is based on the inhibition of ser/thr-proteinphosphatases by Microcystin (MC) congeners. MC congeners are a class of structurally similar cyanobacterial toxins, which are critical to human consumption.

**Conclusion:** We show that it is possible to identify biologically meaningful toxicity signatures by applying the (*α, β*)-*k* feature sets on extended connectivity fingerprint representations of MC congeners. Boolean rules were derived from the feature sets to classify toxicity and can be mapped on the chemical structure, leading to insights on the absence/presence of substructures that can explain toxicity. The presented method can be applied on any other molecular data set and is therefore transferrable to other use cases.

## Background

Methods from Artificial Intelligence and Machine Learning are now increasingly being applied to predict properties of molecules; examples include DeepTox [60], molecular graph attributes [18, 23], ideas from natural language processing [15, 40] and many others [25, 53, 54].

While some of them are achieving impressive performances, they also usually come at the cost of being “black boxes”, meaning that it is difficult for chemists to uncover the structural characteristics behind such phenomenal prediction feats. This situation is not only a characteristic of the Physico-chemical domain on which these methods are applied. They stem from the fact that these methods tend to use all input features encoding the characteristics of a given sample, even though only a few features are often relevant. The situation is not new; for instance, think that even the humble yet powerful linear regression method already has this problem. Unless other restrictions are included to limit the number of variables, this method will produce coefficients for all the input variables. Other powerful methods like those coming from connectionist approaches, such as artificial neural networks (ANNs), use not only all input variables but also include higher-order combinations of them. Untangling which of the input variables are “most important” becomes even more challenging. With the undeniably good performances of the most recent “deep learning” methods based on ANNs, it has been clear that they have their important role in covering. Still, some users would also like some other methods that would naturally lead to “explainable AI” (XAI) techniques that would allow us to go back to the basic characteristics of our samples and generate new knowledge on which are the most important combinations that can explain the outcome of interest [29, 37]. This may later provide useful information in identifying other similar structures in databases and further expand the existing datasets to increase the base of knowledge, which can lead to better experimental designs and chemical elicitation of function from structure. This is also in accordance with the Organisation for Economic Co-operation and Development (OECD) principles for the validation of (quantitative) structure-activity relationship ((Q)SAR) models, which requires (1) defined endpoints, (2) an unambiguous algorithm, (3) a defined domain of applicability, (4) appropriate measures of goodness-of-fit, robustness and predictivity, or (5) a mechanistic interpretation [74]. QSAR tries to relate a set of variables to a specific target or outcome, which is a combinatorial optimization problem [32, 58] and need to be rebuilt for each dataset of interest. In this context, the variables try to encode the chemical structure of a molecule, often resulting in large and sparse matrices [15], which makes it difficult to link to a target outcome [16].

Especially for small dataset, this approach is challenging and prediction quality suffers. There are approaches to circumvent this problem, which includes *e*.*g*. transfer learning. Transfer learning in this context is applying knowledge gained on a large dataset for a similar problem on a smaller dataset to predict molecular properties. These models have been developed to ensure prediction reliability on a small set of compounds, which can be *e*.*g*. congeneric compounds [13, 52]. In addition, it was shown that large numbers of descriptors make it difficult to generalize and interpret resulting QSAR approaches [78] and that feature selection can be used to reduce the number of features (*i*.*e*., low cardinality) to have better model transparency [24, 31].

There are many different methods to encode molecular data that try to capture molecular structure or structural information. In general, different methods can be graph-based (*e*.*g*. [18, 23, 43, 46]), text-based (*e*.*g*. [34, 88, 96]) or numeric (*e*.*g*. [15, 40, 44, 47, 57]) [1]. In this work, we focus on numeric representation with binary characteristics in a sample, *i*.*e*., something that can be either be present or not. We are talking about a ‘predicate’ that, given a sample, it evaluates to either ‘True’ or ‘False’. We will refer to these predicates as “Boolean features” and follow the conventional notation that a ‘1’ indicates that a particular question is answered in the affirmative (True) and a ‘0’ if it is the opposite (False). A well-known example for numerical representation of molecular data are circular fingerprints, *e*.*g*. extended connectivity fingerprint (ECFPs) [85]. In this case, a ‘1’ then would indicate that particular substructure is present in a molecule and a ‘0’ its absence. ECFPs are commonly used to encode molecular structures in cheminformatics [14] to be able to *e*.*g*. compare similarity of molecules [75, 83], or clustering [85], predict molecular properties [60, 97], or discover retrosynthetic routes [87]. It is then clear that, given a set of molecules and molecular descriptions given by ECFPs, the task of identifying a “signature”, that is a subset of molecular descriptions (*i*.*e*., a subset of Boolean Features that can “explain” a particular outcome of interest), is a well-posed combinatorial optimization problem called *k*-Feature Set [21].

Other mixed integer optimization models that have been used to classify molecular data is e.g. the hyper-box framework [98, 101], where multiple hyper-boxes are placed around data points that belong to the same class. With the help of optimisation variables and constraints, multi-class classification problems can be solved and classes predicted. Further developments of this framework even outperformed state-of-the-art classifiers (*i*.*e*., logistic regression, supported vector machines and neural network) [100], and were used to *e*.*g*. generate a rule-based IF-THEN decision model to classify CO_2_ storage sites [93]. In the work of Oooi et al. [73], hyper-box-based machine learning was applied to predict fragrance properties of molecules and deterministic rules were generated to design potential fragrance molecules. The advantage of the hyper-box framework is its interpretability and transparency [73]. Further usage of mathematical optimization is mostly directed towards the design of molecular structure, which is reviewed in Austin et al [6].

In this contribution, we present an approach to predict toxicity by using a method to obtain these signatures that are based on identifying (*α, β*)-*k* feature sets, *i*.*e*., feasible solutions to a generalization of the *k*-Feature Set problem. To achieve this, we use an exact mathematical programming approach that solves an NP-hard problem to optimality. This guarantees that it is quite unique in terms of dimensionality reduction as a pre-processing technique both in the ML and AI spectrum. We note that “machine learning” mainly involves an optimization step of an adaptive system that is addressed in an ad hoc and heuristic manner (and also frequently requiring the fine-tuning of “hyperparameters”). Here we propose a type of exact approach for dimensionality reduction that would not compromise, and in many cases enhance, the generalization properties of a machine learning method used in tandem with this technique.

The dataset used for this study is based on Microcystin (MC) congeners and their ability to inhibit three different ser/thr-proteinphosphatases (PPP): PPP1, PPP2A and PPP5 [2]. They are toxins released during cyanobacterial blooms, which influences drinking water safety worldwide. Several incidences of drinking water closure and human morbidity were reported and linked to MC congeners and cyanobacterial blooms. MC congeners are a class of cyclic heptapeptides with currently over 270 different known congeners [10] which share a common overall structure (Figure 1) [27]. Nevertheless, different congeners have very different capacities to inhibit specific PPPs, and only a few have been tested and analysed [2, 28, 35]. Due to an increasing number of known congeners and difficulty in the ability to synthesize those, *in-silico* approaches are necessary for further investigations and risk assessment to guarantee safety for human consumption [2].

In a first approach by Altaner et al. [2], IC_50_ values (half maximal inhibitory concentration) of 18 MC congeners were determined and a Machine Learning approach was developed to predict those. MC congeners were numerically encoded with Mol2vec [40] and different PPPs with ProtVec [5]. Both encodings are based on Word2vec [62], which is an unsupervised machine learning approach from natural language processing. This results in a vector with fixed dimensions that cannot be tracked back to individual features (or here: molecular or protein substructures). Since performance on this small dataset was relatively poor, synthetic minority oversampling technique was applied to increase the prediction performance by creating new artificial data points for minority classes. Three machine learning models were built: 1) Mol2vec [40] encoding with Random Forests machine learning algorithm, 2) Mol2vec [40] encoding with Extreme Gradient Boosting [17] machine learning algorithm, and 3) Mol2vec [40] and ProtVec [5] encoding with Random Forests machine learning algorithm [12]. The final prediction was based on majority voting. This approach provided 80 to 90 % correct predictions for different toxicity classes and resulted in an overestimation of toxicity, which is not critical in risk assessment but undesired wrong classification. There is only one other approach classifying MC congeners by applying QSAR method with multiple linear regression (MLR). The feature representation chosen in this approach is an average over the whole MC congener structure and does not consider the importance of different substructures. In addition, the median lethal dose (LD50) was the target value of MLR on MC congeners [94].

**Figure 1.**
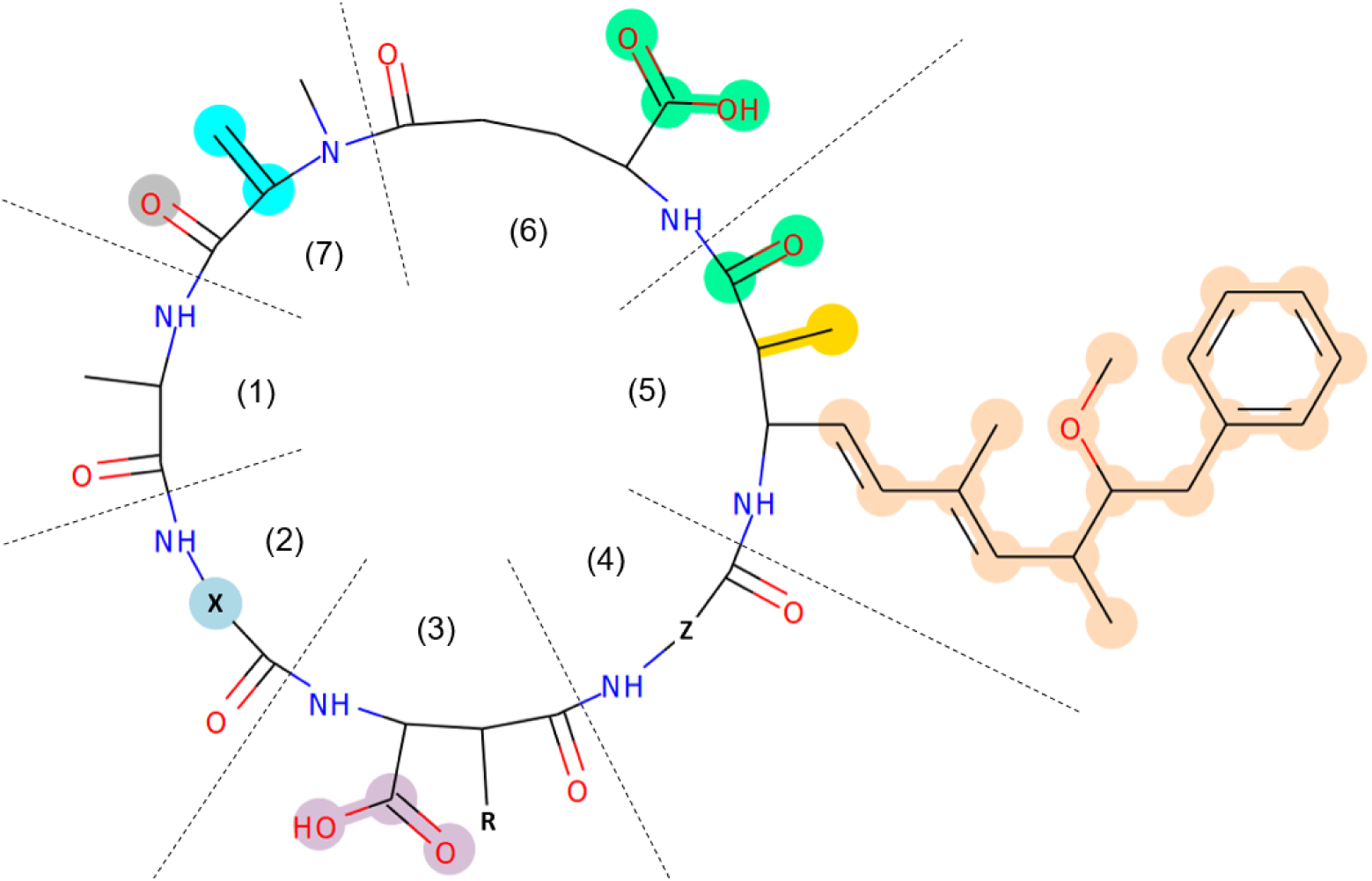
Summary of known interaction sites of MC congeners with PPP1 and PPP2A. X and Z denote different amino acids, R can be a hydrogen or methyl group. The numbers in brackets indicate the residue number in the macrocycle. Different colours indicate different types of interactions: Beige and light blue: hydrophobic interactions; gold: water molecule replacement; mint-green: indirect coordination to metals; cyan: covalent bonding; silver (unique to PPP2A) and light purple: hydrogen bonds. Modified from [27] and [41].

In our work, we show that the identification of (*α, β*)-*k* feature sets (*i*.*e*., exact solutions to a well-defined computational optimization problem) is a straightforward method to be used with ECFPs [85] which provides insights on the structural differences and conserved characteristics that can explain our target of interest (which is identifying substructures related to toxicity in this case). We offer this as a case study of a more general methodology that can be applied to other problems and can also be extended to non-Boolean problems (such as categorically-valued features and classes) or even numerically valued features after suitable approaches for discretization. In contrast to the approach published by Altaner et al. [2] our method explores the encoding of a small molecular dataset with explainable features to be able to identify distinctive structural characteristics of chemical toxicity within a set of structurally similar molecules.

## Methods

### Data curation

The dataset we use has been previously analysed with a machine learning technique [2] so by direct comparison readers can see the advantages of this current method in uncovering the substructures that are associated with toxicity. It contains the IC_50_ values of different but structurally similar molecules on three ser/thr-proteinphosphatases (PPP), namely PPP1, PPP2A, and PPP5. In total, we have 47 samples obtained from these experiments. In the work by Altaner et al. [2], MC congeners were classified in three classes: 1) toxic, 2) less toxic and 3) non-toxic. Since most MC congeners in the dataset belong to the toxic class and only a few to the less toxic and non-toxic class, molecules with IC50 values higher than 40 nM were summarized in the less toxic class. Our final classification is made up of two classes - the toxic and less toxic class.

In our analysis, we have omitted 3 of the 47 samples ([*Enantio*-Adda5]MC-LF for PPP1, PPP2A and PPP5) since they correspond to experiments for which a very large value of IC_50_ was observed, *i*.*e*., the value was considered too high to be measured with the analysis method. Nevertheless, it is reasonable to add those samples to the inactive class (*i*.*e*., less toxic), which can be used later to validate our results. Consequently, we have 17 samples (each corresponding to a different molecule) for both PPP1 and PPP5; for PPP2A we have only 10 samples; see Table 1.

**Table 1.** Distribution of samples per proteins in the dataset.

| protein | # samples | # features |
| --- | --- | --- |
| PPP1 | 17 | 2048 |
| PPP2A | 10 | 2048 |
| PPP5 | 17 | 2048 |

For validation and discussion of the approach, a second dataset was curated for PPP1 by combining the omitted sample from [2] two samples from Fontanillo and Köhn [27] and all available MC congeners with IC_50_ values for PPP1 (in total 5) in Pubchem [48]. This resulted in a dataset with eight additional MC congeners (see Table 2). Both dataset are available in a machine-readable format.

**Table 2.** New IC_50_ values for PPP1. * IC_50_ value is too high to be measured (inactive compound).

| Samp | molecule | IC<sub>50</sub> (nM) | Classification |
| --- | --- | --- | --- |
| 47 | Mol1 [27] | 0.52 | toxic |
| 48 | Mol2 [27] | 253500 | less toxic |
| 49 | 44271302 [68] | 0.8 | toxic |
| 50 | 44271308 [69] | 0.8 | toxic |
| 51 | 44271410 [71] | 0.52 | toxic |
| 52 | 44271411 [72] | 0.3 | toxic |
| 53 | 44271325 [70] | 3.0 | toxic |
| 54 | [*Enantio-Adda5*]MC-LF [2] | * | less toxic |

### Molecular features via extended-connectivity fingerprint

Extended-connectivity fingerprint (ECFP) is one of the most widely used methods to encode molecular data for activity prediction, similarity searches, and many other applications [14]. The ECFP is based on a variant of the Morgan algorithm [63] and a circular fingerprint, encoding substructures of a molecule to capture structural features. It iterates over all atoms of a molecule and encodes the molecule first on an atom level with a radius equal to 0. For each atom, an identifier is generated. The radius is iteratively increased, and an identifier encodes larger molecular substructures. The obtained identifiers are then hashed in an array of fixed size. A value equal to ‘0’ in this fingerprint means the substructure is absent, a value of ‘1’ indicates the substructure is present; thus, the ECFP is a case of a Boolean feature as discussed before. For this contribution, the ECFP4 fingerprint was generated with RDKit with a radius of 2 and a 2048 bit array [50, 85].

### Calculation of **(*α, β*)**-***k***-Feature Set model

We obtained the solution for the instances of the (*α, β*)-*k*-Feature Set model by using an Integer Programming (IP) formulation of the problem as described in [9, 64] and using the commercial software CPLEX to obtain the solutions. The objective function and goal of this combinatorial problem is to minimize the number of features in the solution, with the constraint that we would aim at “covering” the maximum number of *α* nodes (each representing a pair of samples with different target feature), with the constraint that the maximal number of *β*-nodes (each representing a pair of samples with the same target feature) has the same number of features having the same values for those pairs of samples. Such an approach guarantees discrimination while managing within-class similarity [9, 33, 59, 64, 81]. When more than one feasible solution is available for the problem, then the solution providing the greatest coverage of sample pairs belonging to different classes is preferred. Hence, the (*α, β*)-*k*-Feature Set problem offers a minimum set of *k* features, which as a group can well characterize the dataset by maximising the intra-class relationship and inter-classes discrimination information. The IP solution was obtained using the IBM ILOG CPLEX Optimization Studio V12.8.0 (also referred to as CPLEX). The CPLEX software can be downloaded at https://www.ibm.com/support/pages/cplex-optimization-studio-v128.

### Visualisation of molecular features

To visualize the molecular structure and feature sets, RDKit was used to highlight atoms and bonds [50] and Inkscape [39] to arrange images. Due to iteratively increasing the radius around individual atoms for ECFP generation, feature set positions can overlap.

## Theory

### Identification of **(*α, β*)**-***k***-feature sets as signatures of toxicity

In this contribution, we employ an approach based on the (*α, β*)-*k*-Feature Set Problem, a combinatorial problem which is a generalization of the *k*-Feature Set Problem [21]. The decision version of this problem is both NP-Complete and W[2]-Complete [19]. The first result indicates that if the computational complexity classes P and NP are different, then there is no efficient (*i*.*e*., polynomial-time) algorithm for finding exact solutions to this problem. Here P refers to all problems that can be solved in polynomial-time in the size of the instance (input), with NP the class called Non-deterministic polynomial. The second result equally indicates that, according to the current conjectures, there is no fixed-parameter tractable algorithm that can find exact solutions^1^.

While these results seem pessimistic, there are many problems such as this one that is W[2]-Complete, yet some exact solvers can find solutions for large instances in a reasonable amount of time. A similar situation has been observed with this problem. We note that optimization problems associated with NP-complete are generally considered difficult to solve. We have previous experience in bioinformatics applications that indicate that large instances of this problem can be solved to optimality using mathematical programming techniques (e. g. by transforming the original problem and solving an Integer Programming instance using commercial solvers). While we give later a list of previous applications of the approach, it is useful to explain a special case, the *k*-Feature Set.

### The *k*-Feature Set Problem

Using the notation of Davies and Russell [21], the *k*-Feature Set can be formally described as the following decision problem:

*k-* Feature Set

**Instance**: Given a set *X* of examples (which are composed of a binary value specifying the value of the *target feature* and an array of *n* binary values specifying the values of the other features) and a positive integer *k >* 0.

**Question**: Does a set *S* of *k non-target* features exist such that:

- *S* ⊆ {1, · · ·, *n*}, (*i.e*., the set *S* has the indexes of the features in the feature set),
- |*S*| = *k*, (*i.e*., the set *S* has cardinality equal to *k*),
- no two examples in *X* that have identical values for all the features in *S* have different values for the target feature?

### **The** (*α, β*)-*k*-Feature Set and previous applications

The (*α, β*)-*k*-Feature Set Problem is a natural generalization of the *k*-Feature Set Problem and can be defined using the same notation as follows

(*α, β*)-*k*-Feature Set Problem

**Instance:** The same matrix as for the *k*-Feature Set Problem, positive integers *k, α >* 0, and *β* ≥ 0.

**Question**: Does a set *S* of *k non-target* features exist such that:

- *S* ⊆ {1, · · ·, *n*}, (*i*.*e*., the set *S* has the indexes of the features in the feature set),
- |*S*| = *k*, (*i*.*e*., the set *S* has cardinality equal to *k*),
- any two examples in *X* that have different values for the target feature differ in at least the value of *α* features in *S*,
- any two examples in *X* that have the same value for the target feature also have the same value in at least *β* features in *S*?

The (*α, β*)-*k*-Feature Set was first introduced in 2004 in the context of the analysis of microarray datasets [20]. In fact, Cotta, Sloper and Moscato not only introduced this problem in their paper, they also deal with a variant of it in which the optimization task is to find the set of optimal thresholds to binarize the gene expression values such that a (*α, β*)-*k*-feature set with *α* = *β* = *k* can be found. The optimization goal in [20] is to maximize the value of *k*, and an evolutionary algorithm is used to find those thresholds.

Solutions obtained by solving instances the (*α, β*)-*k*-Feature Set Problem have been very useful to derive knowledge from datasets of a wide variety such as political science [65, 66] and business analytics [65], bioinformatics [7], data mining for breast cancer subtype diagnosis [59], cancer cell line subtype identification [51], prediction of clinical symptoms of Alzheimer’s disease [30, 45, 84], microarray data analysis of animal models of Parkinson’s disease [38] and Alzheimer’s disease [7, 64, 80], gene expression signatures in tissue from patients with prostate and breast cancer [42, 61, 79, 81], image analysis [49], and distinguishing childhood absence epilepsy patients from controls by the analysis of their background brain electrical activity [86].

The identification of small *k*-feature sets, a processing method for dimensionality reduction, not only facilitates training in machine learning but also improves generalization [30], even in the case of complex unknown target functions [26]. Specialized heuristics exist to find approximate solutions for large problem instances [22] as well as Integer Programming formulations [8, 9] that allow obtaining exact solutions for problem sizes like the one we are dealing with in this work. Consequently, the solutions reported here are optimal, although they are not guaranteed to be unique since the instance can have many.

To our knowledge, this is the first application of this approach to employ the discovery of (*α, β*)-*k*-Feature Set to datasets generated from ECFP4 and to identify “Boolean signatures of toxicity”. Since the size of our dataset is relatively small in comparison with transcriptomic ones (which involve tens of thousands of features), we can use exact methods such as those of [8, 9] to find the optimal solutions. One possibility is to iterate in the procedure, first identifying one optimal solution and then iteratively solving instances in which one of the features found in the optimal solution is removed. That could lead to sets of optimal solutions.

### An illustrative example of a simple classification scheme based on a single 3-feature set

We illustrate what a feature set is by giving an example of a 3-feature set that we have early observed in our dataset. Table 3 shows a 3-feature set that has been found while separating the samples into two groups: on one side, we have four samples that have a very large value of IC_50_ (samples 14, 15, 16 and 17), corresponding to experiments with [*β*-D-Asp3]-MC-RR, [*β*-D-Asp3, Dhb7]-MC-RR, [Anda5]-MC-LY(Prg) and [Amba5]MC-LY(Prg). All the other samples correspond to the other class.

**Table 3.**
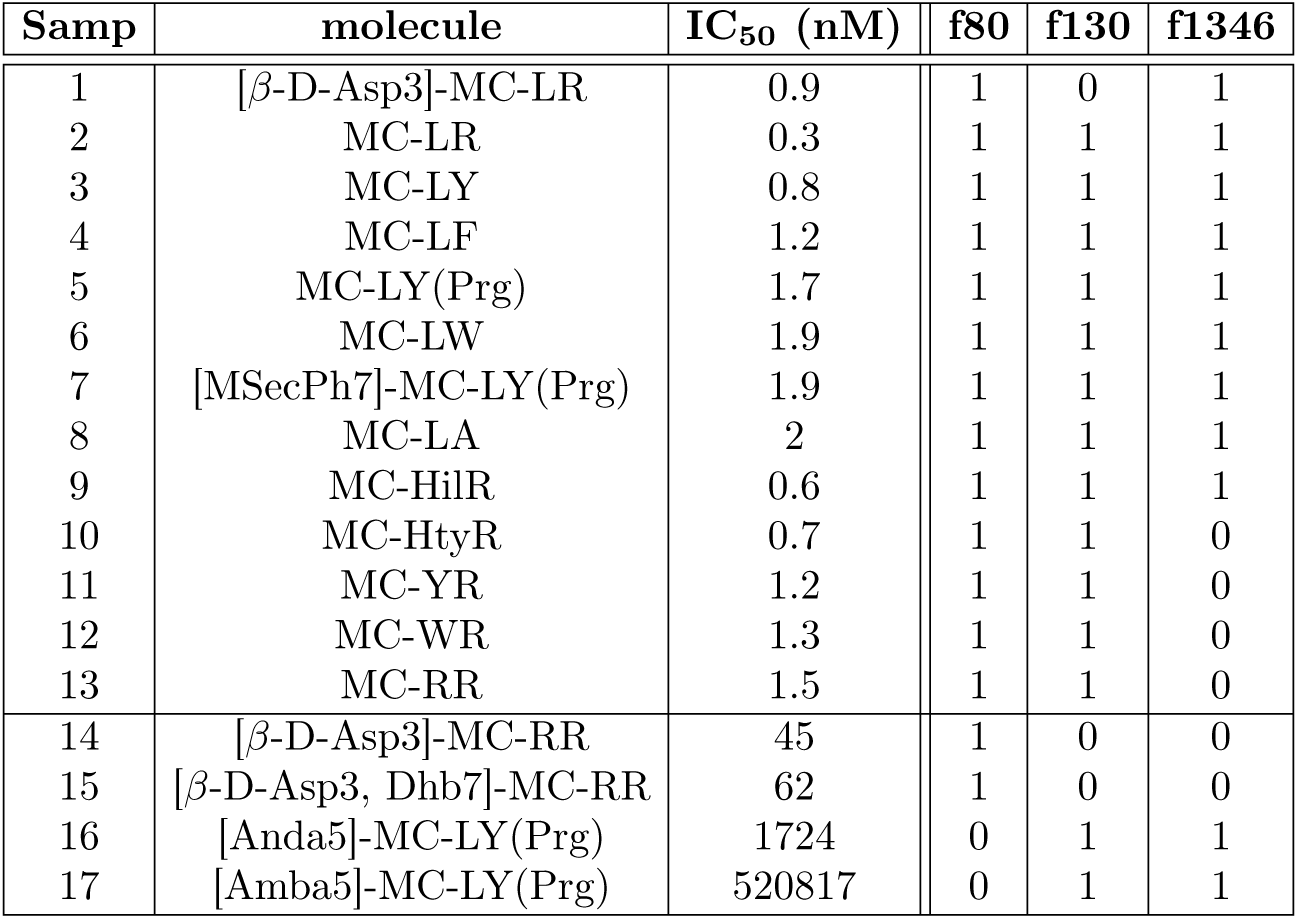
An illustrative example, a 3-feature set, with three ECFP4 substructures, which we observed in the case of PPP1 toxicity.

We can classify the samples into two groups. In the following we will assume that a ‘1’ represents ‘True’ (*i*.*e*., a present substructure), and a ‘0’ represents ‘False’ (or ‘most likely absent’), respectively; then a simple rule arises here: “If either *f* 130 or *f* 1346 is ‘True’ (or both are ‘True’), and in addition *f* 80 is also ‘True’, then it is true that the molecule is toxic for PPP1 (*i*.*e*., it has a low IC_50_ value).” Clearly, for samples 14, 15, 16 and 17 this statement’s premise is not valid. We could also infer another rule: “If (*f* 80 ⊕ (*f* 130 ∨ *f* 1346)) is ‘True’, then the molecule is less toxic for PPP1”, where the symbol ⊕ denotes the exclusive disjunction (the ‘XOR’) and the symbol ∨ is the inclusive disjunction.

These two simple association rules are then able to condense even more the information extracted from the dataset using a *k*-feature set identification approach. We note, however, that this is one possible way of extracting knowledge after a feature set has been obtained. Some researchers may aim at identifying a linear function that “somewhat predicts” the IC_50_ using those three features (via regression methods, for example). Indeed, only two features already do a good job since, for instance, the function *h*(*f* 80*, f* 130) = 3415 − 44 ∗ *f* 130 − 3370 ∗ *f* 80 gives values of IC_50_ of 45 and 3371 for the patterns observed for the samples that are not toxic (14 to 17), a value of 1 for all samples from 2 to 13, and only “misses” what is expected for sample 1 since it also gives the value of 45 (since the expression uses only two of the three features we identified), while the slightly more complex function given by *g*(*f* 80*, f* 130*, f* 1346) = 3415 + 44 ∗ (*f* 130 ∗ *f* 1346 − *f* 130 − *f* 1346) − 3370 ∗ *f* 80 corrects the problem observed with that sample (now returning a value of 1.0 for the IC_50_). Identifying feature sets of small cardinality is an important step to reduce the dimensionality of the machine learning problem and users of the technique can then employ machine learning techniques of their choice, integrating this methodology in a filter approach such as shown in [30, 84].

### Mathematical description of the **(*α, β*)**-***k***-Feature Set Problem

Cotta, Sloper and Moscato introduced the (*α, β*)-*k*-Feature Set Problem as a generalization of the *k*-Feature Set Problem [20] (*i*.*e*., the case in which we set *α* = 0 and *β* = 0 is the NP-complete). This means that the decision version of this combinatorial problem is NP-complete as Davies and Russell already proved in 1994 that *k*-Feature Set is NP-complete [21].

We will use the (*α, β*)-*k*-Feature Set Problem as our mathematical formalization of the task of interest in studying toxicity since we aim to obtain robust “Boolean toxicity signatures” of different molecules, thanks to their fingerprints. This robustness is achieved by finding signatures that likely contain many different feature sets (such as the 3-feature set already discussed) but with a different approach. We aim at having some redundancy of features selected while still allowing for discrimination. Indeed, our Boolean signatures guarantee that “by design”, since we are using an exact algorithm, if a feasible solution exists for the dataset of interest (given our requirements), then at least one *α* ≥ 1 Boolean features will help discriminate between any pair of samples that belong to different classes. In addition, the solution obtained will have at least *β* ≥ 0 similar values between any two samples of the same class. Although we generally search for signatures with *α* ≥ 1 we will formally and in more mathematical detail define the problem as follows:

(*α, β*)-*k*-Feature Set Problem

**Instance**: A set of *m* examples *X* = {*x*^(1)^*, . . ., x*^(^*^m^*^)^}, such that for all 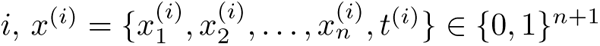, and three integers *k >* 0, and *α, β* ≥ 0.

**Question:** Does there exist an (*α, β*)-*k*-Feature Set 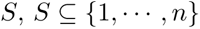 with |*S*| ≤ *k* and such that:

- for all pairs of examples 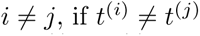 there exists 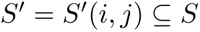 such that 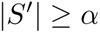 and for all 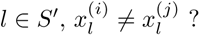
- for all pairs of examples 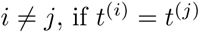 there exists 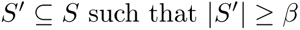 and for all 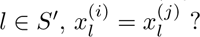

where the set *S^′^* is not the same for all pairs of examples so we have written 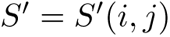.

## Results and discussion

### Identification of Boolean signatures Preprocessing

We first check for any equivalent features where more than one feature has the same set of values for all samples (*i*.*e*., their Hamming distance is zero). We thus reduce the size of the instance by keeping only one feature as a *representative feature* while keeping a record of the others that are equivalent to the one still included). A representative feature is now a feature that represents an entire set of features which are identical in value for all samples. The class label is encoded by the *target feature t*, and the entries are set to ‘1’ for the class of interest and ‘0’ for the others. Those features that have the same value for all samples are called *irrelevant*, and they are also removed during preprocessing.

After the identification of duplicate features from each of the proteins, it remains only:

- 25 representative features in PPP1
- 17 representative features in PPP2A
- 25 representative features in PPP5

### Creation of Boolean signatures on sets of equivalent discriminatory representative features

For each of the three datasets, we run an independent process to obtain a Boolean signature employing the exact solver for the (*α, β*)-*k*-Feature Set Problem. We start by identifying the maximum number of representative features that can discriminate between samples that belong to different classes. For the three datasets, the number found was 2, indicating that for each one of them, there is at least a pair of samples that have only two representative features with different values in each class. While the number is the same, the pair may be different in different datasets. We also note that the representative features currently in the matrix actually are surrogates for all the other equivalent representative features, so this number is higher and is reported after the computation of the signature is completed. We will then denote representative features with capital letters to distinguish them from individual features since we follow standard computer science notation that uses capital letters for sets. For instance, *F* 130, when studying the toxicity of PPP1, represents the set that includes feature *f* 130 and a set of its equivalent features, *i*.*e*., we will write this using set notation as *F* 130 = {*f* 130*, f* 1319*, f* 1780*, f* 1838*, f* 1842}. The list of equivalent features is given in supplementary information section “Equivalent Features of (*α, β*)-*k*-Feature Set Solution”

We then proceed by identifying the minimum value of *k >* 0 for which there exists a feasible solution for the (*α, β*)-*k*-Feature Set Problem with *α* = 2 and *β* ≥ 0. These values have been *k* = 7, *k* = 4 and *k* = 7 once again for the datasets for PPP1, PPP2A and PPP5, respectively. We now know that we are searching for signatures that have *α* = 2 and *k* = 7, so the task is now to find the maximum value of *β* ≥ 0 for which there is still a feasible solution for that instance. These values are 1 for both the cases of PPP1 and PPP5 and 0 for PPP2A. The Boolean signatures are displayed in Table 4.

**Table 4.** Outcome of solving the (*α, β*)-*k*-Feature Set on each of the toxicity discrimination tasks. The solutions obtained are the feasible solutions, such they maximally discriminate between the two classes (*i*.*e*., maximally achievable value for *α*), while they have the maximally achievable value for *β* and minimum value for the total set of features *k*.

| protein | $\alpha$ | $\beta$ | $k$ | Boolean signature of toxicity |
| --- | --- | --- | --- | --- |
| PPP1 | 2 | 1 | 7 | {F32, F130, F232, F295, F336, F695, F1346} |
| PPP2A | 2 | 0 | 4 | {F32, F130, F232, F773} |
| PPP5 | 2 | 1 | 7 | {F32, F130, F232, F295, F336, F773, F1346} |

**Table 5.** Optimal *α* = 2, *β* = 0, and *k* = 4 feature set identified in the case of PPP2A toxicity. Note that a capital ‘F’ indicates that we are referring to a set of equivalent features.

| ID | molecule | IC<sub>50</sub> (nM) | F32 | F130 | F232 | F773 |
| --- | --- | --- | --- | --- | --- | --- |
| 18 | MC-RR | 1.6 | 1 | 1 | 0 | 0 |
| 19 | MC-LR | 0.5 | 1 | 1 | 0 | 0 |
| 20 | MC-YR | 1 | 1 | 1 | 0 | 0 |
| 21 | MC-LW | 0.7 | 1 | 1 | 0 | 0 |
| 22 | MC-LA | 1.4 | 1 | 1 | 0 | 0 |
| 23 | MC-LF | 0.7 | 1 | 1 | 0 | 0 |
| 25 | MC-LY(Prg) | 0.4 | 1 | 1 | 0 | 0 |
| 26 | [MSeCPh7]-MC-LY(Prg) | 0.9 | 1 | 1 | 0 | 0 |
| 24 | [β-D-Asp3, Dhb7]-MC-RR | 84.3 | 1 | 0 | 1 | 0 |
| 28 | [Amba5]-MC-LY(Prg) | 2135 | 0 | 1 | 0 | 1 |

### Toxicity signature for PPP2A

The identified toxicity signatures can then be used for creating simple logical rules to classify samples. Alternatively, other types of classifiers can be used based on the resulting reduced dimensionality datasets. The following is one such simple example of the usage of boolean logic to generate these rules and their structural interpretation.

For PPP2A, and the toxicity signature shown in Table 5, another simple logical rule emerges: *“If either all the features of the sets F232 or all of the features of F773 are ‘True’ (i.e., present in the structure), then the molecule is less toxic for PPP2A”*. From the limited nature of the data, such a rule leaves out some possibility since we have no evidence of what happens when both are true, of course, as no experiment with such an outcome was observed. However, we have another rule that we can infer from the data: *“ ‘If all the features in the set F32 and the set F130 are all ‘True’ (i.e., present in the structure), then the molecule is toxic for PPP2A”*. In fact, given the evidence obtained so far, we should probably restrict the combined use of these rules more by using a conjunction. In natural language, they would then be expressed as: *“If F32 and F130 are both ‘True’ (i.e., present in the structure), and neither F232 or F773 are ‘True’, then the molecule is toxic for PPP2A”*. Analogously we can also reformulate the other rule, *“If either F232 or F773 are ‘True’ (i.e., present in the structure) and it is not the case that both F32 and F130 are present in the structure, then it the molecule is less toxic for PPP2A”*.

Figure 2 shows how we can identify the structures represented by the signatures. The use of other types of classifiers may generate different methods to predict activity. This brief example indicates how the proposed method can generate knowledge about toxicity drivers. To estimate their biological importance, identified feature sets were compared to data summarized in a review by Fontanillo and Köhn [27]. Figure 2a shows feature sets F32 and F232 on [*β*-D-Asp3, Dhb7]-MC-RR, while Figure 2b shows feature sets F130 and F773 on [Amba5]-MC-LY(Prg). Both MC congeners are less toxic, as we can infer from the IC_50_ values and Boolean rules. In comparison to the known functional groups of MC congeners (Figure 1), the feature sets determined here are coarser and include a lot more atoms compared to the detailed structure-activity-relationship study presented by Fontanillo and Köhn. This result is expected since we were only able to include 10 MC congeners to determine a Boolean signature of toxicity. The feature set F32 is mainly made up of the Adda residue, which is important for binding towards PPP1 and PPP2A but not toxic itself [67, 95]. In comparison, if Adda is missing, almost no activity towards PPP2A can be observed [11]. In detail, if there is a methyl group (which is true for molecules with F773) instead of the Adda residue, submicromolar activity can be observed, which is classified as essentially non-toxic. This is in agreement with the IC_50_ values of our dataset and F773. In Figure 2b we can see that the Adda residue is missing, and F773 is present instead of F32. F32 and F773 also include a methyl group, which is important to block active site water (Figure 1, gold colour).

**Figure 2.**
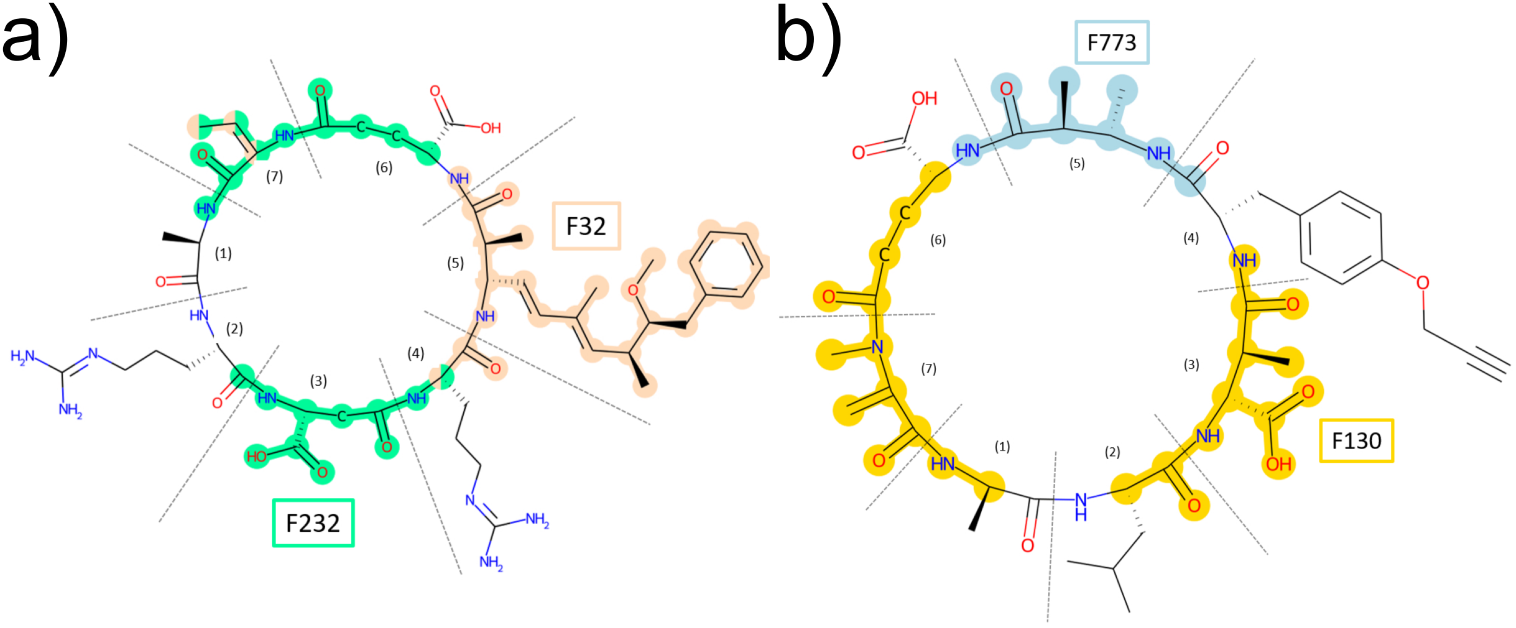
Feature set of PPP2A highlighted on three example structures. F32 is beige and contributes to toxicity. F773 is light blue and Adda is replaced by a methyl group, which leads to decreased toxicity. The role of the carboxyl group at position 3 in F130 (gold) and F232 (mint-green) is not fully understood. The methyl group in F130 (gold) might block the interaction. The covalent bond at position 7 in F232 (mint-green) is not possible due to modification, but not essential for toxicity. a) [*β*-D-Asp3, Dhb7]-MC-RR and b) [Amba5]-MC-LY(Prg).

F232 and F130 are widespread across the backbone of the MC structure. Both include three important functional groups for hydrogen bond-building (Figure 1, silver and light purple colour) and the covalent bond (Figure 1, cyan colour). Interestingly, the hydrogen bond of the carboxylic oxygen in position seven is specific for interaction with PPP2A and found in both F232 and F130, but not in the fingerprint sets for PPP1 and PPP5. This indicates that there is indeed biological meaning in our fingerprint sets. It is known in literature that carboxylic acid of *β*-D-MeAsp in position three builds a hydrogen bond with PPP2A and PPP1, but its role is not fully understood and investigated yet [36, 77]. Covalent bonding of MC congeners to PPPs is an important mechanism to irreversible bind to PPPs, but does not affect toxicity, since MC congeners also bind reversible to PPPs, [55, 56]. Hence, it is not further considered.

When we compare our evaluation to the Boolean rules, we define a molecule as toxic if F32 and F130 are true and F232 and F773 are false. This is in agreement with known interactions in literature [27]. F773 indicates the Adda residue is missing (and replaced by a methyl group), which results in inactivity. When F232 and F130 are compared, the only difference is the group responsible for covalent bonding. Since a methyl group is added in F232, it might be able to block the nearby hydrogen bond of the carbonyl group and thereby to influence toxicity and binding.

### Toxicity signature for PPP1

**Table 6.**
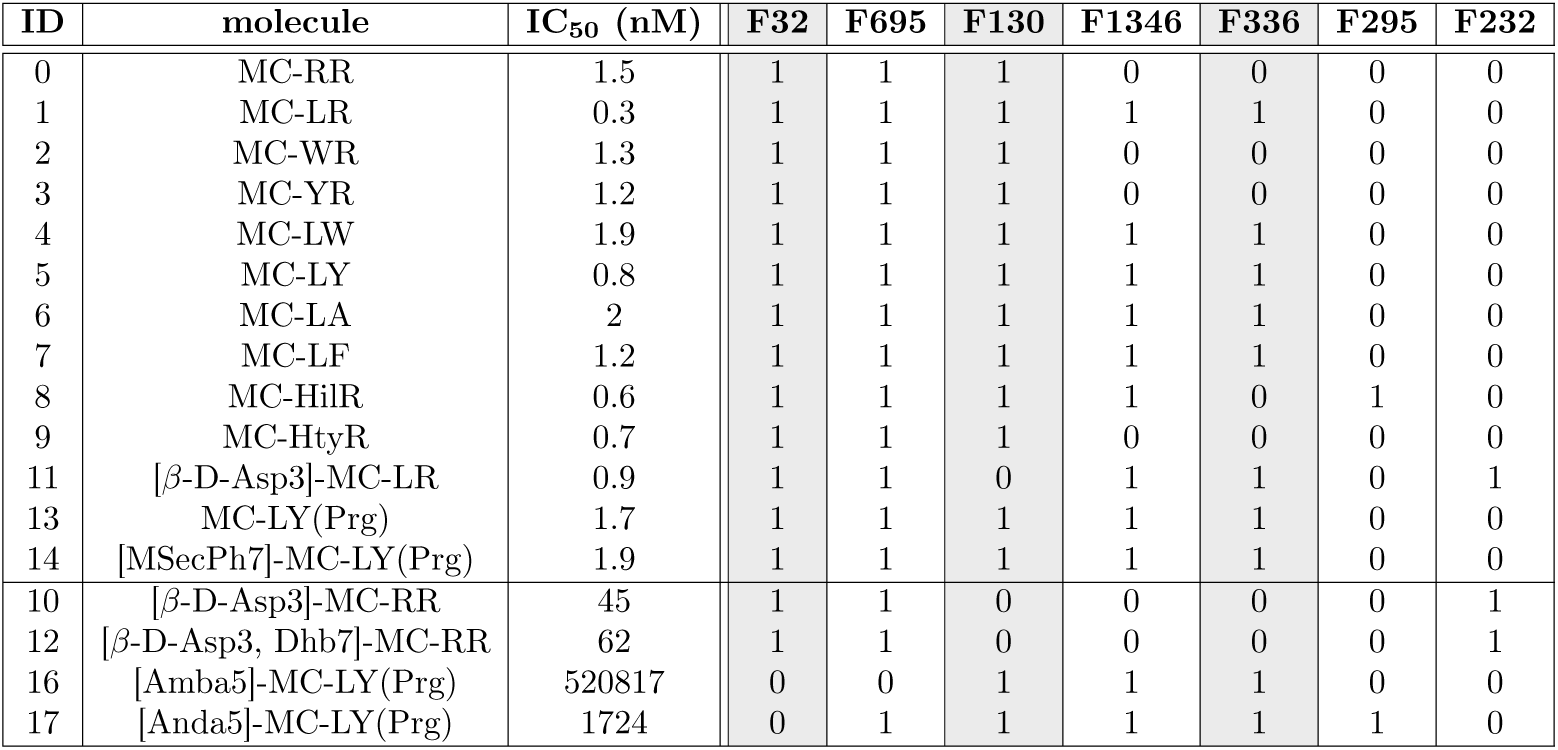
Optimal *α* = 2, *k* = 7, *β* = 1 feature set which was found in the case of PPP1 toxicity. Note that a capital ‘F’ indicates that we are referring to a set of equivalent features.

For PPP1 we can infer from Table 6: *“If either all the features in the signature F130 or all the features in the signature of F336 are ‘True’ (i.e., all the features are present in the structure in isolation or both are) and also all the features of the signature of F32 are ‘True’, then the molecule is toxic for PPP1; otherwise, it is less toxic.”* Interestingly, for the three feature sets involved in this rule, and from all the possible patterns that we can have (*i*.*e*., 8 = 2^3^ for three Boolean features sets in this case), we have seven patterns out of the eight. The only missing pattern is when all the three features sets are all ‘False’, and no such a sample exists in the original database.

For PPP1 the feature sets were also overlaid on two example structures covering all feature sets (Figure 3). To estimate their biological importance, identified feature sets were compared to data summarized in a review by Fontanillo and Köhn (Figure 1) [27]. Figure 3a shows feature sets F32, F695, F1346, F336 and F232, while Figure 3b shows feature sets F32, F695, F130, F1346 and F295. Both MC congeners are toxic, as we can infer from the IC_50_ values in Table 6 and the Boolean rule. In contrast to the feature sets for PPP2A, a more fine-grained distinction between the feature sets are visible.

Nevertheless, they are still more coarse in comparison to the detailed structure-activity-relationship study presented by Fontanillo and Köhn [27]. The feature set F32 is also for PPP1 covering the side chain of the Adda residue, which was already discussed in the previous section for PPP2A. It is important for binding towards PPP1 and PPP2A, but not toxic itself [67, 95]. Feature sets F130 (gold) and F232 (mint-green) (Figure 3) either represent *β*-D-Me-Asp or *β*-D-Asp as residue 3. The carboxylic group of residue 3 is important for hydrogen bonding with PPP1 and PPP2A, but as mentioned earlier, the role is not fully understood. Also, studies showed different results for demethylation of *β*-D-Me-Asp. Some showed negative effects for binding towards PPP1, others showed that there is no effect [36, 89]. Here, we almost have identical IC_50_ values for both molecules towards PPP1: 0.90 nM for [*β*-D-Asp3]MC-LR versus 0.30 nM for MC-LR. Nevertheless, when we compare MC-RR (1.5 nM) to [*β*-D-Asp3]-MC-RR (45 nM), high differences in IC_50_ values can be observed, and demethylation causes less toxicity towards PPP1. Accordingly, F130 (gold) is the methylated variation and contributes to toxicity, as correctly identified by our method. In our Boolean rule, it is captured that either F130 (gold) or F336 (silver) needs to be present in addition to F32 for a compound to be toxic. Since [*β*-D-Asp3]MC-LR is demethylated but still toxic, we assume that F336 (silver) is important for toxicity. This observation can also be found in literature: it is known that a hydrophobic residue at position 2 and hydrophilic at position 4 (which is not recognized here and also not highlighted in the structure-activity relationship study) is important for higher inhibitory potency [82]. F336 (silver) corresponds to a hydrophobic side chain (here leucine) which is important for binding. When we again compare to [*β*-D-Asp3]-MC-RR, a basic amino acid (arginine) is present at this position and exhibits lower toxicity compared to [*β*-D-Asp3]MC-LR. Since MC-RR still has high inhibitory potency towards PPP1, we conclude that both a basic amino acid at position 2 and demethylation of *β*-D-MeAsp3 is necessary to have lower inhibitory potency. In contrast to our feature sets for PPP2A, the atom responsible for covalent binding is not identified in a feature set here, which is uncritical, since covalent bonding is not required for toxicity [56] and not a key process for inhibition [55].

**Figure 3.**
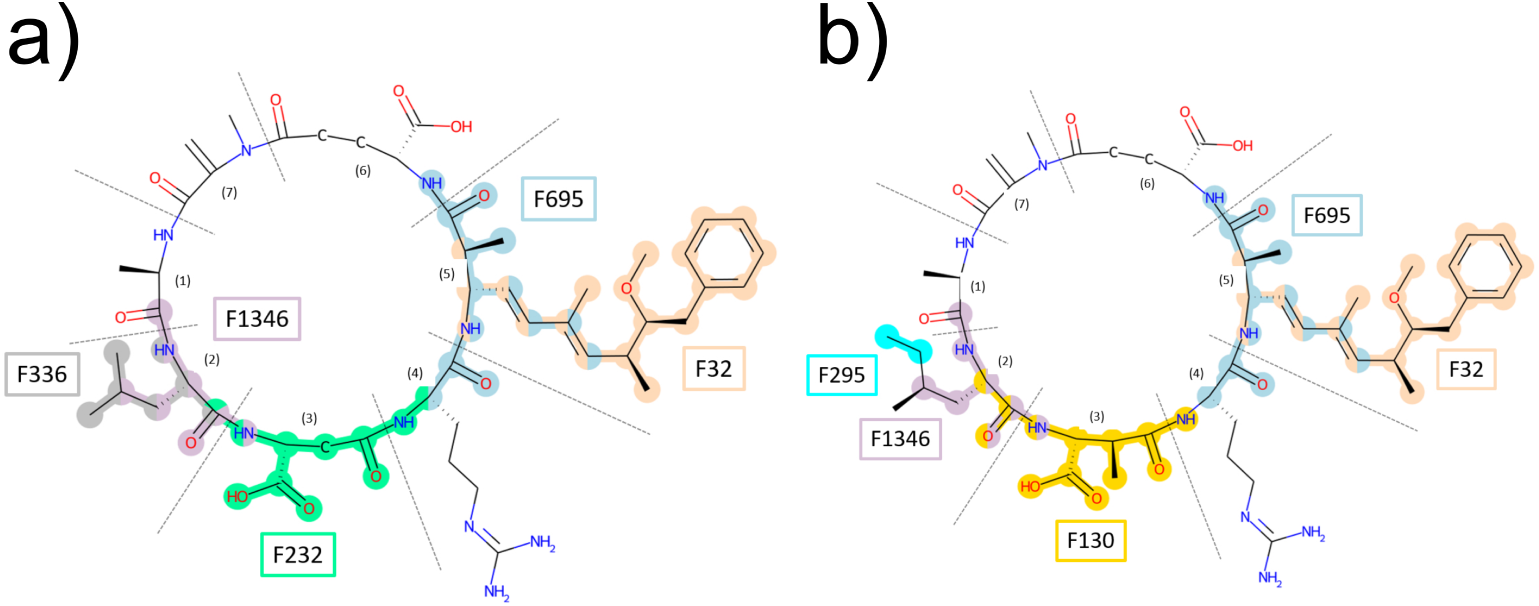
Feature set of PPP1 highlighted on two example structures. F32 is beige and marks the Adda residue, which is important for binding and therefore toxicity. The role of F130 (gold) and F232 (mint-green) is not fully understood and studies showed different results. In the dataset used for our study demethylation causes reduced toxicity towards PPP1. F336 (silver) is known to contribute to toxicity, as a hydrophobic residue at position 2 increases toxicity. This residue is also partly covered by F1346 (light purple) and F295 (cyan). F695 (light blue) captures water replacement and coordination to metal ions, which contributes to binding and therefore toxicity. a) [*β*-D-Asp3]MC-LR and b) MC-HilR.

### Toxicity signature for PPP5

For PPP5, we also have seven sets of features in the signature of toxicity, sharing many with PPP1 (six, see Table 7).

In this case three sets help to separate the samples in two classes and the only missing pattern is the one in which all the three features are not present, which no sample exists in the original database, neither for PPP1 nor for PPP5.

The same rule inferred for toxicity for PPP1 is also valid for PPP5, so we can now update our previous rule as: *“If either f336 or f130 are ‘True’ (i.e., present in the structure in isolation or both are) and also f32 is present, then the molecule is toxic for both PPP1 and PPP5; otherwise it is less toxic.”*

PP5 is less studied than PPP1 and PPP2A; for this reason, less data is available to compare to literature. It is known that the three-dimensional structure of the catalytic core matches well with PPP1 [99] and PPP2A, as the root mean square deviation of pairwise /*α*-carbon is between 1.3 and 1.5 Angström [91, 92]. Nevertheless, there are differences in sequence and structure. When calculating percent identity by aligning the protein sequence of the catalytic unit of PPP5 in comparison to PPP1 and PPP2A with Protein Blast, [3] a percent identity of 44.40 % and 41.02 % was calculated, respectively. So PPP5 is indeed more closely related to PPP1 than PPP2A. Despite, PPP1 and PPP2A share a higher percent identity of 49.27 %.

**Table 7.** Optimal *α* = 2, *k* = 7, and *β* = 1 feature set solution which was found in the case of PPP5 toxicity. Note that a capital ‘F’ indicates that we are referring to a set of equivalent features.

| ID | molecule | IC<sub>50</sub> (nM) | F32 | F130 | F1346 | F336 | F295 | F232 | F773 |
| --- | --- | --- | --- | --- | --- | --- | --- | --- | --- |
| 29 | MC-RR | 11.7 | 1 | 1 | 0 | 0 | 0 | 0 | 0 |
| 30 | MC-LR | 5.1 | 1 | 1 | 1 | 1 | 0 | 0 | 0 |
| 31 | MC-WR | 5.1 | 1 | 1 | 0 | 0 | 0 | 0 | 0 |
| 32 | MC-YR | 5.6 | 1 | 1 | 0 | 0 | 0 | 0 | 0 |
| 33 | MC-LW | 6.1 | 1 | 1 | 1 | 1 | 0 | 0 | 0 |
| 34 | MC-LY | 4.1 | 1 | 1 | 1 | 1 | 0 | 0 | 0 |
| 35 | MC-LA | 4.7 | 1 | 1 | 1 | 1 | 0 | 0 | 0 |
| 36 | MC-LF | 2.5 | 1 | 1 | 1 | 1 | 0 | 0 | 0 |
| 37 | MC-HilR | 4.2 | 1 | 1 | 1 | 0 | 1 | 0 | 0 |
| 38 | MC-HtyR | 4.7 | 1 | 1 | 0 | 0 | 0 | 0 | 0 |
| 40 | [β-D-Asp3]-MC-LR | 10.2 | 1 | 0 | 1 | 1 | 0 | 1 | 0 |
| 42 | MC-LY(Prg) | 1.7 | 1 | 1 | 1 | 1 | 0 | 0 | 0 |
| 43 | [MSecPh7]-MC-LY(Prg) | 18.2 | 1 | 1 | 1 | 1 | 0 | 0 | 0 |
| 39 | [β-D-Asp3]-MC-RR | 167.1 | 1 | 0 | 0 | 0 | 0 | 1 | 0 |
| 41 | [β-D-Asp3, Dhb7]-MC-RR | 877.1 | 1 | 0 | 0 | 0 | 0 | 1 | 0 |
| 45 | [Amba5]-MC-LY(Prg) | 54063 | 0 | 1 | 1 | 1 | 0 | 0 | 1 |
| 46 | [Anda5]-MC-LY(Prg) | 2420 | 0 | 1 | 1 | 1 | 1 | 0 | 0 |

**Figure 4.**
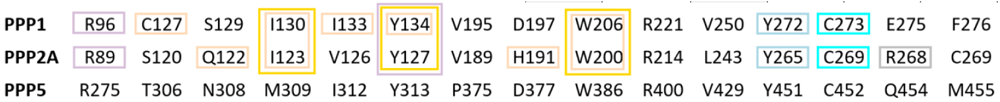
Partial sequence alignment of PPP1, PPP2A and PPP5. Amino acids involved in interactions with MC congeners [27] are marked. Different colours indicate different interactions: beige and light blue: hydrophobic interactions; gold: water molecule replacement; mint-green: indirect coordination to metals; cyan: covalent bonding; silver (unique to PPP2A) and light purple: hydrogen bonds. The partial sequence alignment is modified from Pereira et al [76].

**Figure 5.**
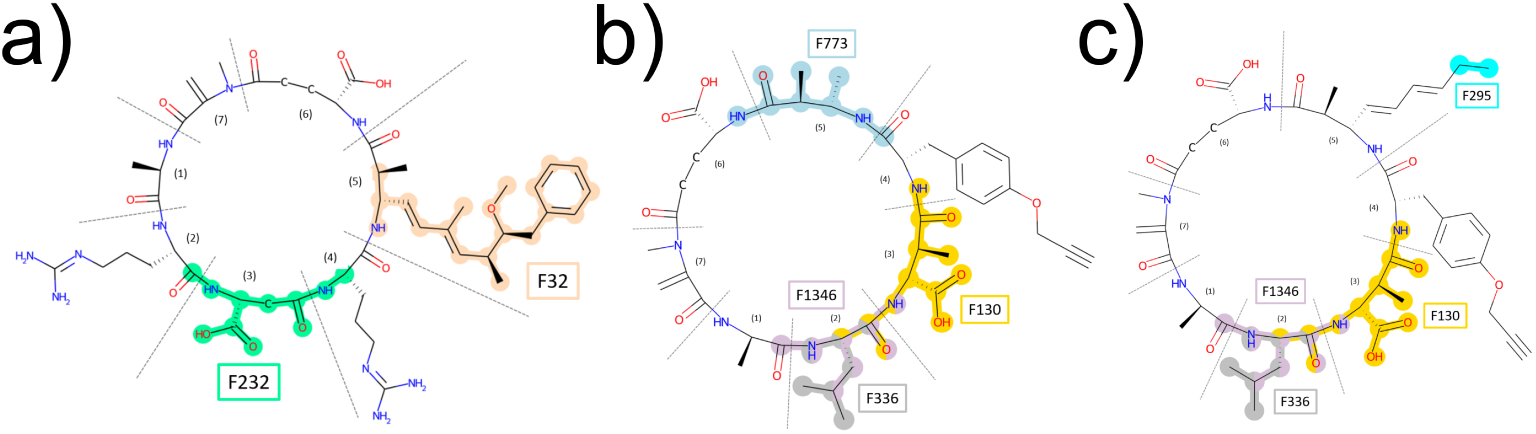
Feature set of PPP5 highlighted on three example structures. F32 is beige and marks the Adda residue, which is important for binding. If it is missing, binding is reduced, which is shown by F773 (light blue). The role of F295 (cyan) is unclear, but toxicity is highly reduced compared to F32. The role of F130 (gold) and F232 (mint green) is unclear. F336 (silver) and F1346 (light purple) mark a hydrophobic residue which contributes to toxicity at position 2. a) [*β*-D-Asp3]MC-RR, b) [Amba5]-MC-LY(Prg), and c) [Anda5]-MC-LY(Prg).

Structure-Activity Relationship studies for PPP5 are rare compared to PPP1 and PPP2A and mostly consider metal coordination in the binding site, catalytic mechanism or other toxins than MC congeners. This has to be taken into consideration to understand the relevance of our feature sets to understand how useful they are for PPP5. We could find a partial amino acid sequence alignment of PPP1, PPP2A and PPP5 in literature (Figure 4). From this alignment, we concluded which residues might be important for interaction and mapped them to interactions found with PPP1 and PPP2A. Of course, this is only a first step on an approach suggested by our findings and further investigations will be needed, as for example, follow-up mutational studies are necessary to provide more evidence. Nevertheless, the sequence alignment revealed that we have similar interactions as for PPP1 and PPP2A, since amino acids were either identical or had similar chemical properties.

We discussed that PPP5 shares a percent identity of over 40 % with PPP1 and PPP2A and that PPP1 and PPP2A share a percent identity of 49.27 %. Accordingly, feature sets obtained for different PPPs overlap, and also individual features can overlay due to the feature generation process with ECFP4. The feature sets were overlaid on three example structures covering all feature sets (Figure 5). Apart from the sequence alignment, we can not conclude on the interaction between PPP5 and MC congeners. Consequently, we refer the reader to the previous sections’ discussion of feature sets for PPP1 and PPP2A.

### Considering new structures

Results obtained by solving to optimality the (*α, β*)-*k*-Feature Set Problem are going to be dependent on the dataset used for a specific problem. As we mentioned before, an augmented dataset may have different, perhaps more restricted solutions to the problem (with smaller values of the maximum values of *α*, *β*, and perhaps also a different value of *k* for which a feasible solution can be found).

For this reason, a second dataset was built from left out or literature values to discuss the selected feature set for PPP1 and the stability of the Boolean rule, see also Table 8. The (*α, β*)-*k*-Feature Set is constructed from identical results of the ECFP4, *i*.*e*., all fingerprint positions have the same values (either 0 or 1). In the case of the new dataset, we obtain different values for the individual fingerprint positions. For this reason, some feature sets had to be separated to achieve distinction into different classes.

In case the fingerprint positions have different values, the feature sets were split up, but the original naming scheme was kept for clarity. The feature sets which are important for toxicity of PPP1 are shown in Table 8 and the list of equivalent features is given in supplementary information section “Equivalent Features of (*α, β*)-*k*-Feature Set Solution”.

From the original dataset, the rule inferred for PPP1 toxicity is the following: *“If either all the features in the signature F130 or all the features in the signature of F336 are ‘True’ (i.e., all the features are present in the structure in isolation or both are) and also all the features of the signature of F32 are ‘True’, then the molecule is toxic for PPP1; otherwise it is less toxic.”* Here, F32 splits up into two feature sets, F32a and F32b, which includes 17 and 3 fingerprint positions, respectively. Since F32a is made up of most of the original fingerprint positions, the original rule was modified as follows: *“If either all the features in the signature F130 or all the features in the signature of F336 are ‘True’ (i.e., all the features are present in the structure in isolation or both are) and also all the features of the signature of F32a are ‘True’, then the molecule is toxic for PPP1; otherwise it is less toxic.”* The new feature sets are shown in Figure 6. When applying the modified boolean rule derived from dataset 1 on dataset 2, we can classify the MC congeners as shown in Table 8. Of a total of 8 new molecules, 6 were classified correctly with 5 toxic and 1 less toxic molecule, which is a total of 75 %. The remaining two molecules classified incorrectly are 44271410 and [*Enantio*-Adda5]MC-LF. 44271410 is a toxic molecule classified incorrectly as less toxic, since F130 and F336 are both 0. [*Enantio*-Adda5]MC-LF is a less toxic stereoisomer of the toxic MC-LF. Only four stereocenters are flipped, which is not captured in the ECFP4. For this reason, it is classified incorrectly, as the same ECFP4 already describes a toxic molecule.

This analysis shows that we can use the Boolean rule also for classifying new compounds and have a reliable estimate of their toxicity. Nevertheless, previous work has shown that using structural alerts on molecular structures can be problematic. In the work of Stepan et al. [90], it was shown that for a high percentage of approved drugs, one or more alerts in their structure could be identified, even though no significant incidences were reported. This leads to the assumption that structural alerts may over-exaggerate toxicity of molecules which can be problematic in drug discovery, but not for toxins, [90] which is the focus of this work. Alves et al. [4], discuss that specific structural features, as they are used in structural alerts, do not act individually but rather act together with other various substructures, which do not need to be in close proximity. These various substructures do define the biological effect in the end, which questions the usage of fragments to predict molecular toxicity of the whole molecule [4]. Our approach presented here does focus on substructures, but instead of using individual substructures, multiple of them are summarised in feature sets, representing several substructures instead of individual fragments. In addition, the boolean rule defined to aid in determining relevant structures for toxicity contribution is considering multiple feature sets, and therefore does not focus on one substructure at one position of the molecule. For this reason, we think that applying the (*α, β*)-*k* feature set problem to the MC congener dataset is a valid approach to identify substructures leading to toxicity of MC congeners. A reminder note, transferability is limited, as we have only a very small test set of 17 molecules and 2 out of 8 incorrectly classified new molecules. However, if more data becomes available, retraining would be beneficial to benefit from the extra samples and possibly lead to more accurate results. Such expanded dataset is currently not available, so we present here *a first approach* to predict MC congener toxicity, the dimensionality of the dataset and the size of the molecules allow us to understand the benefit of uncovering substructures.

**Table 8.** IC_50_ class, predicted class and split up feature set solution for new dataset on PPP1 toxicity. Each column has a capital letter actually representing a set of features that are equivalent as described below.

| ID | molecule | IC<sub>50</sub> Class | Predicted Class | F32a | F32b | F130 | F232a | F232b | F232c | F232d | F295 | F336 | F695 | F1346a | F1346b |
| --- | --- | --- | --- | --- | --- | --- | --- | --- | --- | --- | --- | --- | --- | --- | --- |
| 47 | Mol1 [27] | toxic | toxic | 1 | 1 | 1 | 0 | 0 | 0 | 0 | 0 | 0 | 1 | 0 | 0 |
| 48 | Mol2 [27] | less toxic | less toxic | 0 | 1 | 0 | 1 | 1 | 1 | 1 | 0 | 1 | 0 | 1 | 1 |
| 49 | 44271302 [68] | toxic | toxic | 1 | 1 | 0 | 1 | 0 | 1 | 1 | 1 | 1 | 1 | 0 | 1 |
| 50 | 44271308 [69] | toxic | toxic | 1 | 1 | 0 | 1 | 0 | 0 | 1 | 0 | 1 | 1 | 0 | 1 |
| 51 | 44271410 [71] | toxic | less toxic | 1 | 1 | 0 | 1 | 0 | 0 | 0 | 0 | 0 | 1 | 0 | 0 |
| 52 | 44271411 [72] | toxic | toxic | 1 | 1 | 0 | 1 | 0 | 0 | 0 | 0 | 1 | 1 | 0 | 1 |
| 53 | 44271325 [70] | toxic | toxic | 1 | 1 | 0 | 1 | 0 | 0 | 0 | 0 | 1 | 1 | 0 | 1 |
| 54 | [Enantio-Adda5] MC-LF [2] | less toxic | toxic | 1 | 1 | 1 | 0 | 0 | 0 | 0 | 0 | 1 | 1 | 1 | 1 |

## Conclusion

In this work, we presented an approach to obtain meaningful toxicity signatures of molecules by identifying (*α, β*)-*k* feature sets. We could show that the identification of (*α, β*)-*k* feature sets can be directly applied on ECFP4 to provide novel insights on chemical structure, in particular to identify molecular substructures related to toxicity. The fingerprint sets built by the (*α, β*)-*k* method were used to derive Boolean rules to determine toxicity on PPP1, PPP2A and PPP5. Data from literature was collected and compared to the fingerprint sets to investigate whether there is biological meaning in the fingerprint sets. We could show that there is indeed biological meaning in the fingerprint sets that can explain MC congener toxicity towards PPPs. Since there are more than 270 MC congeners described in the literature (but not tested), in a second step, the method was applied to a new dataset collected from literature to estimate the performance on molecules with unknown toxicity towards PPP1. A dataset of 8 MC congeners was collected, of which most congeners could be classified correctly. This result shows that the feature sets and Boolean rules are also applicable to new dataset. In this work, we dealt with a difficult problem in which the number of examples used for training (17 compounds) was two orders of magnitude smaller than the number of samples, yet the rules derived from simple Boolean rules were able to correctly classify 75 % of the samples in the test set. There are several directions for future work, including using an ensemble of classifiers for this unbalanced dataset [33], investigating a larger set of MC congeners when new data is available. In addition, studying cases with larger molecules and representations that include ECFP4 with stereocenters will further improve results and lead to a more general applicability in the future.

**Figure 6.**
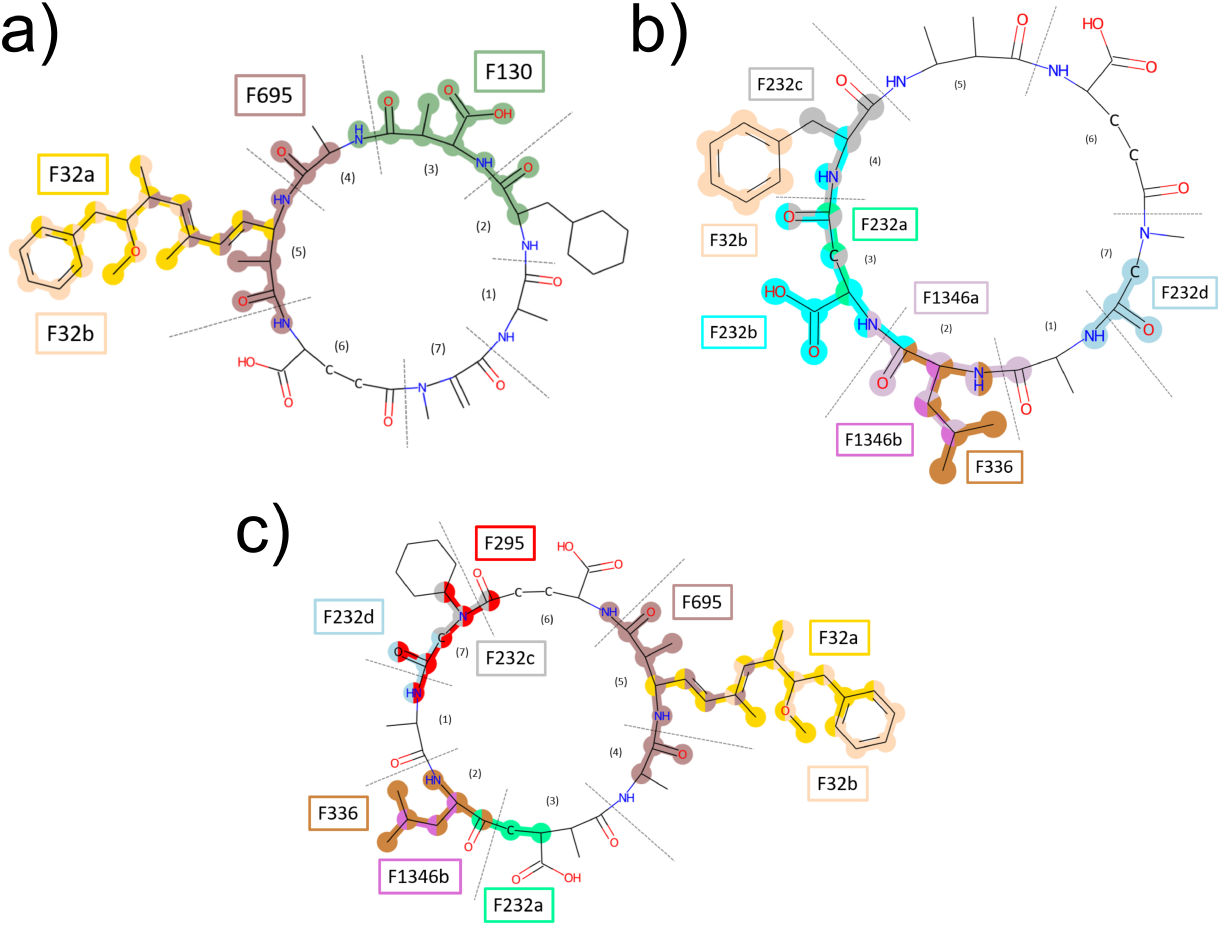
New feature set of PPP1 highlighted on three example structures. The colour coding is as follows: FP32a is gold, FP32b is beige, FP130 dark green, FP232a is mint-green, FP232b is cyan, FP232c is silver, FP232d is light blue, FP295 is red, FP336 is brown, FP695 dark brown, FP1346a is light purple, and FP1346b is purple. a) Mol1, Mol2, c) 44271302.

## Supporting Information

- Additional File 1: PDF: List of equivalent features sets and respective ECFP4 indices.

## Funding

This research was supported by the Australian Government through the previous support from Australian Research Council’s Future Fellowship (FT120100060) and Discovery Projects funding scheme (projects DP120102576, DP140104183), and the current Discovery Projects funding scheme (project DP200102364). P.M. acknowledges a generous donation from the Maitland Cancer Appeal. This work was partly funded by the Deutsche Forschungsgemeinschaft (DFG, German Research Foundation) – Project-ID 251654672 – TRR 161.

## Acknowledgements

PM and MNH thank Prof. Regina Berretta for discussions about how to generate signatures using Integer Programming models. SJH and FS thank Daniel R. Dietrich for fruitful discussions and gratefully acknowledge the Konstanz Research School of Chemical Biology (KoRS-CB).

## Footnotes

1 https://en.wikipedia.org/wiki/Parameterized_complexity

## Notes

### Competing Interest Statement

The authors have declared no competing interest.

