## Additional File 1 for "The (*α, β*)-*k* Boolean Signatures of Molecular Toxicity: Microcystin as a Case Study"

### Supplementary Information: The $(\alpha, \beta)$-$k$ Boolean Signatures of Molecular Toxicity: Microcystin as a Case Study

Pablo Moscato[1,*], Sabrina Jaeger-Honz[2,*], Mohammad Nazmul Haque[1], Falk Schreiber[2,3],

**1 School of Information and Physical Sciences, College of Engineering, Science and Environment, University of Newcastle, Newcastle, Australia**
**2 Department of Computer and Information Science, University of Konstanz, Konstanz, Germany**
**3 Faculty of Information Technology, Monash University, Clayton, Australia**
*****

#### Structural considerations and equivalent features

As we have discussed in the manuscript, each feature corresponds to a different substructure within the molecule. When the absence and presence of two substructures are the same for all the samples in the dataset, they are deemed equivalent. While unfortunately some machine learning algorithms, like the so-called *wrappers* for instance, would select either one or the other in the quest of finding "explanations" with a small number of features, this leads to some loss of information. In contrast, our procedure, by keeping a "bag" of equivalent features, does not suffer from this issue and the reduction of dimensionality (by just considering one representative) should be always followed by an expansion, "unpacking" the information we have condensed in each set of equivalent individual features.

It is then entirely possible that at this stage either problem domain knowledge (or an expanded/augmented dataset) may clarify which are the most important features of this "bag", or alternatively, still consider all of them for further investigation. We now discuss what is the case for toxicity of these three proteins since the study case is small yet complex enough to illustrate this post-processing final step to uncover useful knowledge.

##### Equivalent Features of $(\alpha, \beta)$-$k$-Feature Set Solution

###### PPP1

The Boolean signature of toxicity for PPP1 contains the following surrogate features: {F32, F130, F232, F295, F336, F695, F1346}. The feature $f32$, for instance, is equivalent to f80 and many others (when restricted to the samples tested against PPP1). Here we list the equivalent features (if any) for each of those features. We represent the name of the set (i. e. the surrogate feature) capitalizing the name of the first listed feature in the set:

- $F32 = \{f32, f80, f144, f614, f746, f842, f858, f890, f898, f976, f1018, f1067, f1089, f1179, f1200, f1258, f1352, f1520, f1563, f1743\}$
- $F130 = \{f130, f1319, f1780, f1838, f1842\}$
- $F232 = \{f232, f546, f685, f1412, f1434, f1576\}$

- $F295 = \{f295\}$ (i. e. there is no other equivalent feature in this case)
- $F336 = \{f336, f1848, f1877\}$
- $F695 = \{f695, f716, f734, f1125, f1229, f2026\}$
- $F1346 = \{f1346, f1719\}$

##### PPP2A

The Boolean signature of toxicity for PPP2A contains following surrogate features: {F32, F130, F232, F773}. Here we list the equivalent features (if any) for each of those features.

- $F32 = \{f32, f80, f144, f614, f695, f716, f734, f746, f842, f858, f890, f898, f976, f1018, f1067, f1089, f1125, f1179, f1200, f1229, f1258, f1352, f1520, f1563, f1743, f2026\}$
- $F130 = \{f130, f799, f936, f1001, f1146, f1319, f1780, f1838, f1842, f2009\}$
- $F232 = \{f232, f546, f685, f747, f845, f906, f969, f1061, f1070, f1175, f1256, f1412, f1434, f1576, f1822, f1998\}$
- $F773 = \{f773, f1301, f1554, f1604\}$

##### PPP5

Finally, the Boolean $(\alpha, \beta)$-$k$-Feature Set signature of toxicity for PPP5 contains the following surrogate features: {F32, F130, F232, F295, F336, F773, F1346}. Here we list the equivalent features (if any) for each of those features.

- $F32 = \{f32, f80, f144, f614, f746, f842, f858, f890, f898, f976, f1018, f1067, f1089, f1179, f1200, f1258, f1352, f1520, f1563, f1743\}$
- $F130 = \{f130, f1319, f1780, f1838, f1842\}$
- $F232 = \{f232, f546, f685, f1412, f1434, f1576\}$
- $F295 = \{f295\}$ (i. e. there is no other equivalent feature in this case)
- $F336 = \{f336, f1848, f1877\}$
- $F773 = \{f773, f1301, f1554, f1604\}$
- $F1346 = \{f1346, f1719\}$

##### PPP1 and data set 2

In the case of new molecules (or a new data set) the feature sets will be separated due to newly available information. The new feature set obtained after analysing data set 2 is as follows:

- $F32a = \{f32, f80, f144, f614, f746, f842, f858, f890, f898, f976, f1018, f1179, f1258, f1352, f1520, f1563, f1743\}$
- $F32b = \{f1067, f1089, f1200\}$
- $F130 = \{f130, f1319, f1780, f1838, f1842\}$
- $F232a = \{f232\}$ (i. e. there is no other equivalent feature in this case)

- $F232b = \{f546, f1412, f1434\}$
- $F232c = \{f685\}$ (i. e. there is no other equivalent feature in this case)
- $F232d = \{f1576\}$ (i. e. there is no other equivalent feature in this case)
- $F295 = \{f295\}$ (i. e. there is no other equivalent feature in this case)
- $F336 = \{f336, f1848, f1877\}$
- $F695 = \{f695, f716, f734, f1125, f1229, f2026\}$
- $F1346a = \{f1346\}$ (i. e. there is no other equivalent feature in this case)
- $F1346b = \{f1719\}$ (i. e. there is no other equivalent feature in this case)
